# ECM deposition is driven by caveolin1-dependent regulation of exosomal biogenesis and cargo sorting

**DOI:** 10.1101/405506

**Authors:** Lucas Albacete-Albacete, Inmaculada Navarro-Lérida, Juan Antonio Lopez, Inés Martín-Padura, Alma M. Astudillo, Michael Van-Der-Heyden, Jesús Balsinde, Gertraud Orend, Jesús Vazquez, Miguel Ángel del Pozo

**Affiliations:** Mechanoadaptation & Caveolae Biology Laboratory; Cell and Developmental Biology Area, Centro Nacional de Investigaciones Cardiovasculares Carlos III (CNIC). Melchor Fernández Almagro, 3, 28029, Madrid, Spain; Cardiovascular Proteomics Unit. Centro Nacional de Investigaciones Cardiovasculares Carlos III (CNIC), Melchor Fernández Almagro, 3, 28029, Madrid, Spain; Instituto de Biología y Genética Molecular, Consejo Superior de Investigaciones Científicas, Universidad de Valladolid, 47003, Valladolid, Spain, and Centro de Investigación Biomédica en Red de Diabetes y Enfermedades Metabólicas Asociadas, Instituto de Salud Carlos III, 28029, Madrid, Spain; INSERM U1109 - MN3T, The Microenvironmental Niche in Tumorigenesis and Targeted Therapy, 3 avenue Molière, 67200 Strasbourg, Université de Strasbourg, LabEx Medalis, Fédération de Médecine Translationnelle de Strasbourg (FMTS), France

## Abstract

The composition and physical properties of the extracellular matrix (ECM) critically influence tumour cell behaviour, and ECM deposition and remodelling by stromal fibroblast populations is therefore pivotal for tumour progression. The molecular mechanisms by which stromal and tumour cell populations regulate ECM layering are poorly understood. Tumour-stroma interaction is critically dependent on cell-cell communication mediated by exosomes, small vesicles secreted by most cell types and generated within multivesicular bodies (MVBs). Here, we show that caveolin-1 (Cav1), an essential regulator of stromal remodelling and tumour cell fate, plays a central role in modulating both exosome biogenesis and exosomal protein cargo sorting through cholesterol-dependent mechanisms. Quantitative proteomics profiling revealed that a major share of Cav1-dependent exosomal cargoes are compsed of ECM proteins, one of the most important components being tenascin-C (TnC). Comparative functional assays demonstrated that Cav1 is required for fibroblast-derived exosomes to depose ECM and promote tumour cell invasiveness. Exosomes purified from Cav1WT cells, but not those from Cav1-null cells, were able to nucleate distant stromal niches in different organs *in vivo*. These findings suggest a key role for Cav1 as a cholesterol rheostat in MVBs, and seems to determine ECM deposition by eliciting ECM component sorting into specific exosome pools. These results, together with previous work, support a model in which Cav1 is a central regulatory hub for tumour-stroma interactions through a novel exosome-dependent ECM deposition mechanism.

## INTRODUCTION

The tumour microenvironment is increasingly recognized as a key factor in multiple stages of disease progression, particularly local invasion and distant metastasis^1^-^3^. Cancer-associated fibroblasts (CAFs) are an important cell population in different carcinomas and constitute a major source of paracrine growth factors and ECM components for the tumour stroma ^4^,^5^. However, understanding is limited on the molecular mechanisms that determine fibroblast ECM deposition. Recent studies confirm that ECM deposition ultimately depends on a functional core secretion machinery^6^, but further mechanistic characterization of ECM deposition is needed to fully characterize the broad impact of the ECM in health and disease and explore new therapeutic opportunities.

An important mechanism governing tumour-stroma interaction is exosomal cell-cell communication. Exosomes are secreted vesicles containing a variety of cargoes, and are generated from intraluminal vesicles (ILVs) in subpopulations of multivesicular bodies (MVBs), which are membrane-bound compartments intimately linked to the endosomal system^7^-^9^. Secreted by most cell types, exosomes enable a given cell to modulate the behaviour of other cell populations through the transfer of effector or regulatory molecules over long (inter-organ) distances^10^,^11^. Recent studies indicate that tumour-derived exosomes can promote pre-metastatic niche formation^12^-^14^. However, it remains unclear whether and how fibroblast-derived exosomes might modulate these phenomena during tumour progression.

Exosomal cargo composition is heterogeneous and complex, containing a variety of proteins, small ligand peptides and regulatory nucleic acids^9^. It is clear that exosome composition, established by the donor cell, defines the modulatory effect on the receptor population, but little is known about how cargoes are selectively sorted into exosomes. Current evidence supports the co-existence of mechanisms dependent on the endosomal sorting complexes required for transport (ESCRT) machinery with ESCRT-independent mechanisms^15^,^16^, likely reflecting the heterogeneity of MVBs and exosomes and their potential differential impact on health and disease.

Exosome biogenesis is also modulated by lipids through their ability to laterally segregate into specific membrane subdomains, which allows sorting platforms to be organized on the limiting membrane of MVBs ^17^. Cholesterol is likely to be a highly relevant lipid in this context, given its significantly higher abundance in MVBs than in other endomembrane compartments ^18^. Cholesterol is necessary for the formation of high-curvature membrane structures such as caveolae and synaptic vesicles, and a role in the intraluminal sorting of membranes destined for exosome secretion has been also proposed^19^-^21^. However, information remains scarce on the mechanisms modulating cholesterol levels in MVBs and their impact on exosome biology.

Contact sites between endosomes/MVBs and the ER membrane have recently been demonstrated as sites for the specific transfer of proteins and lipids (especially cholesterol). Modulation of ER-MVB contacts depends on the interaction between different proteins, especially members of the ORP family (ORP1L and ORP5). Cholesterol accumulation in MVBs has been reported to reduce ER-MVB contacts ^22^.

Endomembrane trafficking is regulated by caveolin-1 (Cav1), an essential scaffolding component of invaginated plasmalemal nanodomains (*caveolae*). Cav1 participates in several essential cell functions, including sensing and transduction of mechanical cues, endocytosis, spatiotemporal control of signal transduction, and cholesterol homeostasis. Moreover, we recently identified Cav1 as an integral component of mitochondria-associated ER membranes (MAMs), which are specialized ER regions that allow communication with mitochondria ^23^. The pleiotropic functions of Cav1 support a pervasive yet context-dependent role in cancer progression and metastasis ^24^. These diverse Cav1 functions include biomechanical ^25^ and compositional ^26^ remodelling of the ECM by CAFs to favour tumour invasion and metastasis, but the mechanisms through which Cav1 regulates ECM composition are unknown. Cav1 has been detected in exosomes^27^, but it has not been determined whether exosomal secretion is a specific means for Cav1-dependent ECM remodelling.

Here we demonstrate that Cav1 is a constitutive component of exosomes produced by several cell types, including primary CAFs. We show that ubiquitination-acceptor lysines at the N-terminal region of Cav1 are essential for its sorting into the MVB compartment. Comparative proteomic and lipidomic studies showed that Cav1 modulates the sorting of exosome cargo and the segregation of exosome subpopulations through a cholesterol-dependent mechanism. Intriguingly, exosomes derived from Cav1WT fibroblasts are significantly enriched in ECM components, including TnC, an important driver of tumour progression. Complementary *in vitro* approaches indicate that exosomal biogenesis and release are essential for fibroblast ECM deposition. Supporting Cav1-dependence of exosome competency for ECM deposition, exosomes derived from Cav1-null fibroblasts failed to promote protrusiveness, motility and invasiveness in a breast tumour cell line. Cav1- dependent remote ECM deposition by exosomes was recapitulated *in vivo*, suggesting an important role for stromal Cav1 in distant premetastatic niche formation in addition to its reported role in local ECM remodelling. We propose that Cav1 is a key regulatory hub promoting and coordinating primary and metastatic stromal environments by modulating both exosome biogenesis and specific exosomal cargo sorting. This regulation is exerted through the modulation of MVB membrane plasticity, which depends on cholesterol content and organization.

## RESULTS

### Internalized Cav1 redistributes to LE/MVB to its sorting in exosomes

Caveolin-1 internalization is tightly regulated^28^-^30^. Although endocytosed Cav1 has commonly been regarded as committed to degradation^31^,^32^, alternative fates cannot be excluded^33^. To better define this process, we conducted confocal microscopy analysis of endogenous Cav1 trafficking dynamics in mouse embryonic fibroblasts (MEFs) and COS7 cells upon treatment with sodium orthovanadate, which induces Cav1 internalization by protecting Cav1 phosphorylation at tyrosine 14^30^. Orthovanadate exposure induced Cav1 phosphorylation, which correlated with increased colocalization with the late endosome (LE; immunostained with a monoclonal antibody to lyso-bis-phosphatidic acid (LBPA)) and multivesicular bodies (MVBs; immunodecorated for the CD63 marker) (Fig. 1a and Supplementary Fig.1a). Similarly, a significant fraction of Cav1 relocalized to the LE/MVB compartment upon exposure of cells to EDTA, which induces Cav1 endocytosis by promoting cell detachment ^34^, and in response to RNAi-mediated depletion of PTRF, an essencial caveolar coat component required for caveolae stabilization^35^ (Supplementary Fig. 1b and 1c). Interestingly, PTRF depletion induced significant Cav1 phosphorylation (Supplementary Fig. 1c). High-resolution confocal microscopy confirmed the constitutive presence of Cav1 in the MVB lumen, colocalizing with the exosomal marker CD63 in normal conditions (Fig. 1b). To confirm Cav1 internalization in LE and MVB lumens, we generated enlarged endosomes by overexpressing a constitutively active Rab5 mutant (Q79L)^36^. Discrete patches of Cav1 clearly localized to ILVs within MVBs (Fig. 1c), delineated by accumulation of the specific exosomal markers Tsg101 and CD63 (Supplementary Fig. 1d). These observations demonstrate that MVBs are one of the destinies of endocytosed Cav1.

**Figure 1.**
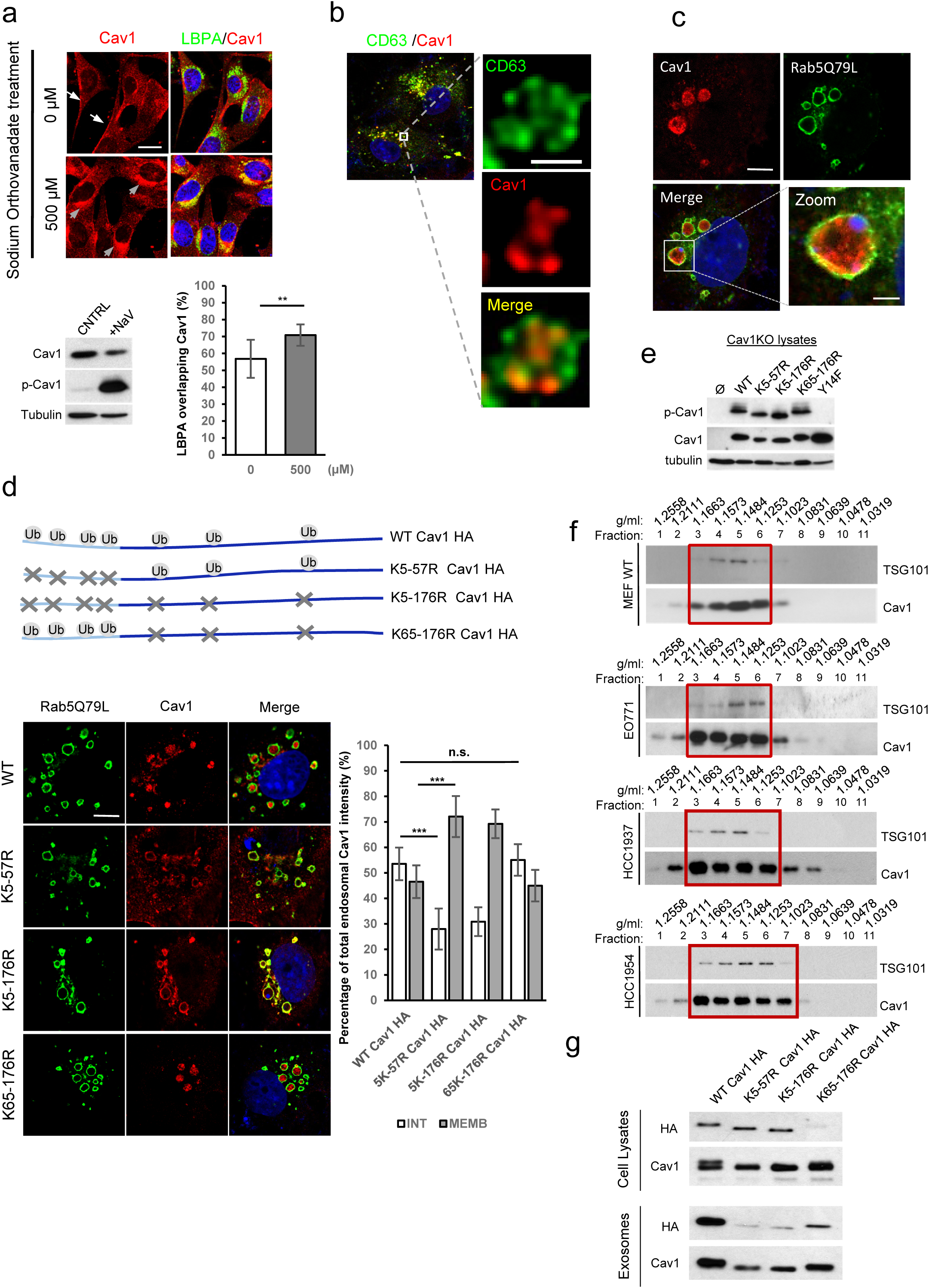
Internalized Cav1 is sorted to exosomes (a) Confocal microscopy of caveolin-1 (red) and LBPA (green) in Cav1WT MEFs after treatment with sodium orthovanadate for 2 h. White arrows indicate plasma membrane Cav1 localization and grey arrows mark Cav1 relocalization to the perinuclear area (Scale bar, 25 µm). Charts show quantification of Cav1-LBPA colocalization. Error bars represent mean±s.d; n=80 cells. Representative western blot of phospho-Cav1 and total Cav1 in sodium orthovanadate-treated cells. (b) Colocalization between Cav1 and the MVB/exosome marker CD63. The right panels show high-resolution views of an MVB compartment after image deconvolution (Scale bar, 1 µm). (c) Cav1 distribution in Rab5(Q79L)-COS7-expressing cells (Scale bar, 10 µm). A zoomed view of an endosome is shown on the right (Scale bar, 2,5µm). (d) *Top.* Ubiquitination-target lysine residues (Ub) in WT Cav1 and engineered versions with the indicated residues mutated to arginine (X). *Bottom.* Confocal analysis of COS7 cells transfected with Rab5(Q79L) (green) and the indicated Cav1 proteins. Ubiquitination is lost upon mutation of the N-terminal region lysines (K5-57R) or all lysines (K5-176R), but not lysines 65-176 (K65-176R) (as reported in Kirchner et al., 2012) (Scale bar, 10 µm). The chart compares localization of Cav1 variants in the endosomal lumen (internal, INT: white) versus the endosomal limiting membrane (peripheral, MEMB: grey), expressed as a percentage of total endosomal localization. Error bars are mean±s.d; n>20 endosomes. (e) Western blot analysis of sodium orthovanadate-induced phosphorylation in WT Cav1, ubiquitination-target lysine mutants, and non-phosphorylatable Y14F Cav1. (f) Cav1 inclusion in exosomes derived from fibroblasts (MEFs). Representative western blots of exosomal proteins floated on a continuous sucrose gradient (0.25-2M). Individual 1ml fractions were collected and after protein precipitation were loaded on electrophoresis gels and analyzed for Cav1 and the exosome marker Tsg101. (g) Western blot analysis of Cav1 in cell lysates and exosomes of COS-7 cells expressing HA-tagged Cav1 ubiquitination mutants.

Phosphorylation and ubiquitination engage in an intricate interplay, with phosphorylation often serving as a trigger for subsequent ubiquitination of adjacent acceptor residues^37^. Ubiquitination and deubiquitination appear to be important steps for protein sorting into ILVs within MVBs. To assess the importance of ubiquitination for Cav1 internalization into MVBs, we analyzed the subcellular distribution of Cav1 mutants in which arginine residues replace lysine residues that act as ubiquitination acceptors (Fig. 1d, upper panel). All mutant forms nicely decorated the outer enlarged-Rab5 endosome membrane/MVB; however, mutation of the N-terminal lysine cluster (K5-57R and K5-176R mutants) abolished Cav1 sorting into ILVs (Fig. 1d). Mutation of Cav1 N-terminal lysines hasbeen shown to impair Cav1 ubiquitination ^38^. This result correlated with lower overall colocalization of these mutants with CD63 (Supplementary Fig. 1e). In contrast, mutation of lysine residues in the carboxyl-terminal Cav1 region (K65-176R mutant) did not affect the ability of Cav1 proteins to localize in MVBs (Fig. 1d). Interestingly, the overall phosphorylation level in these Cav1 mutants was not significantly altered (Fig. 1e), reinforcing the notion of a two-step mechanism driving Cav1 sorting into MVBs, with phosphorylation first eliciting Cav1 internalization towards the outer MVB membrane and ubiquitination then acting as a check-point for proper Cav1 sorting into the ILVs.

To test whether the ILV-localized Cav1 pool is effectively sorted into exosomes, we analyzed density gradient-purified exosomes from different cell lines (including tumour and fibroblast cells) by western blotting. Endogenous Cav1 mainly partitioned with Tsg101, confirming that exosomes can carry Cav1 (Fig. 1f). Interestingly, Cav1 sorting to exosomes was favoured in PTRF-depleted MEFs (Supplmentary Figure 1f), opposite to the impaired sorting of Cav1 mutants with disrupted N-terminal ubiquitination (Fig 1g). Taken together, these data strongly suggest that a significant proportion of internalized Cav1 sorts to the MBV lumen in an ubiquitin-dependent manner, favouring its incorporation into exosomes. Furthermore, internalization of endogenous Cav1 into exosomes is a constitutive process in all Cav1- expressing cell lines tested.

### Cav1 regulates exosomal biogenesis through modulation of MVB cholesterol content

A variety of mechanisms can modulate the composition and organization of membranes incorporated into MVBs, resulting in different ILV and exosome populations^9^. Because Cav1 is a key organizer of membrane architecture across cell compartments^24^, we explored whether Cav1 plays an active role in exosome biogenesis. Genetic ablation of Cav1 did not trigger a general, non-specific alteration of exosomal biogenesis; indeed, Cav1KO fibroblasts had even higher rates of exosome secretion, as determined by western blot of total purified exosomes from equal cell numbers (Fig. 2a upper panel and Figure 2b), sedimentation across sucrose density gradients (Fig. 2a lower panel), and physical characterization with Nanosight technology and transmission electron microscopy (TEM) (Fig. 2c and Fig. 2d)^39^. However, Nanosight and TEM identified two differently sized exosome subpopulations in Cav1WT fibroblasts, whereas Cav1KO fibroblasts produced a homogeneous population containing the smaller exosomes (Fig. 2c and 2d), further suggesting a significant role for Cav1 in exosome biogenesis.

**Figure 2.**
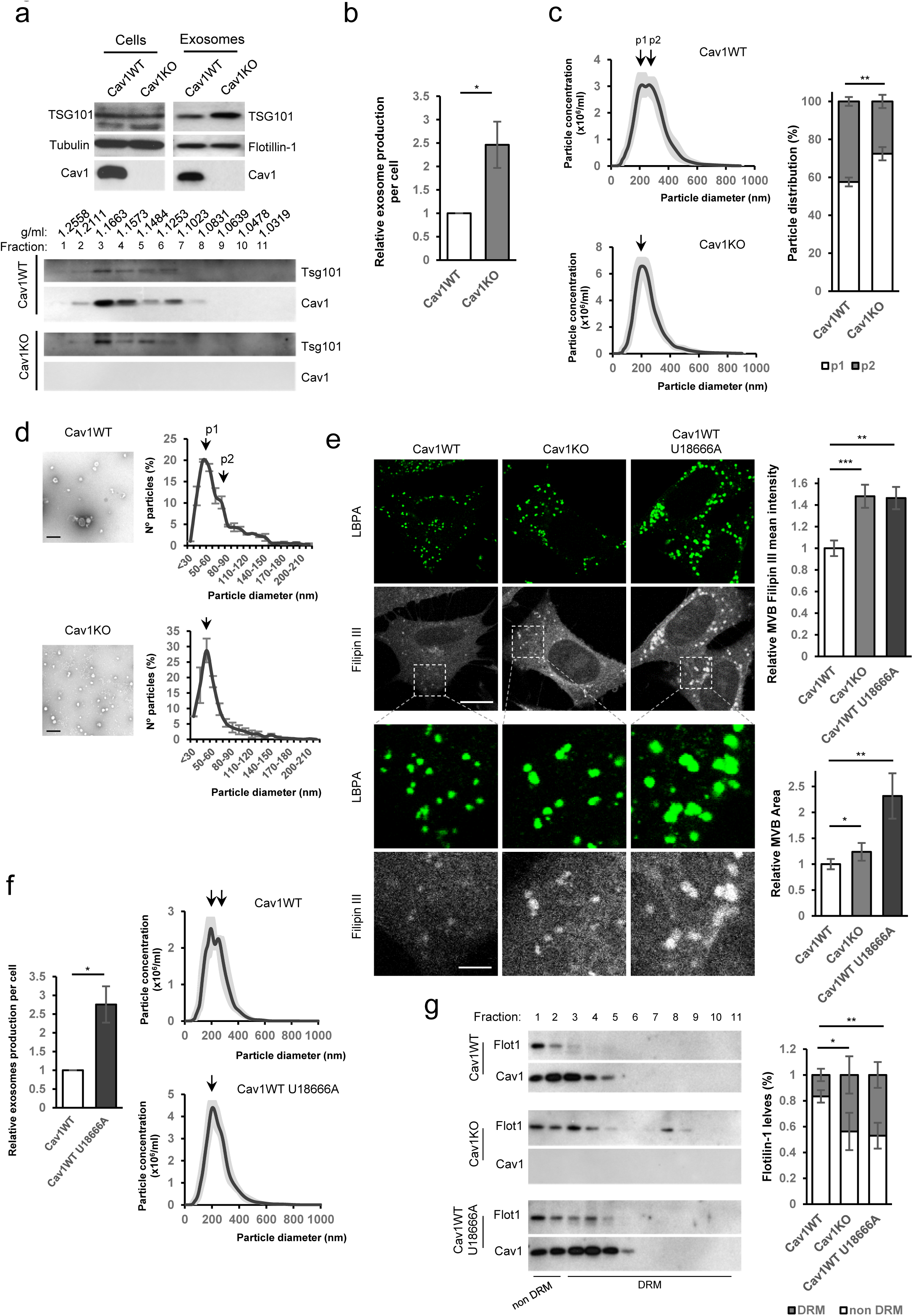
Cav1 regulates exosome biogenesis by modulating MVB cholesterol content Exosomes were isolated from supernatants of Cav1WT and Cav1KO fibroblast cultures. (a) *Top.* Western blots of Cav1, flotillin and the exosome marker Tsg101 in exosome extracts derived from the same number of cells. *Bottom.* Western blots showing the distribution of Cav1 and Tsg101 in exosomes floated on a continuous sucrose gradient. (b) Quantification of exosome particles relative to producing cell number. (c) Nanosight distribution profiles of exosome preparations from Cav1WT and Cav1KO fibroblasts, showing greater morphological and size heterogeneity of Cav1WT-derived exosomes. The right panel shows quantification of the 2 exosome populations identified in Cav1WT and Cav1KO fibroblasts. Error bars are mean±s.e.m; n=10. (d) Representative transmission electron micrographs (TEM) of Cav1WT and Cav1KO exosomes (Scale bar 200 nm). Distribution profiles measured from TEMs of Cav1WT and Cav1KO fibroblasts-derived exosomes. (e) Filipin staining (in grey) and LBPA (in green) of Cav1WT, Cav1KO and U18666A-treated Cav1WT fibroblasts (Scale bar, 20 µm). The lower panel rows show zoomed views (Scale bar, 6 µm). Charts on the right show quantitative analysis of filipin mean fluorescence intensity in MVBs (upper) and total MVB area (lower). (f) Exosome production per cell (left) and Nanosight distribution profiles of exosome preparations (right) from Cav1WT fibroblasts without treatment (control) and treated with U18666A. (g) Western blots of sucrose density gradient fractions in the presence of triton X- 100 of exosomes produced by untreated Cav1WT and Cav1KO fibroblasts and U1866A- treated Cav1WT fibroblasts. The chart shows the proportion of flotillin in detergent-resistant and non-resistant membranes. Error bars are mean±s.d; n=3.

The ability of Cav1 to associate with specific lipids, particularly cholesterol, and to generate specialized membrane organization domains ^24^,^40^prompted us to study whether Cav1 promotes the exosome heterogeneity observed in preparations purified from Cav1WT MEFs by modulating cholesterol levels and membrane organization in MVBs. To explore this, Cav1WT and Cav1KO MEF cells were labelled with filipin, a well-known cholesterol marker. While the number of LBPA-positive structures did not differ between Cav1WT and Cav1KO fibroblasts, MVB area and filipin intensity (a proxy for mean cholesterol content) were significantly higher in Cav1KO cells (Figure 2e). This phenotype was recapitulated by siRNA depletion of Cav1 in Cav1WT cells (Supplementary Fig. 2a). Interestingly, treatment of Cav1WT MEFs with U1866A, a specific promotor of cholesterol accumulation in the LE/MVB compartment^41^, increased exosome secretion while reducing exosome heterogeneity, thus mimmicking the phenotype of Cav1KO cells (Fig. 2f).

A major share of MVB cholesterol is contained in ILVs^18^. We confirmed this cholesterol accumulation in enlarged endosomes by overexpressing constitutively active Rab5Q79L (Supplementary Figure 2b). To confirm whether altered cholesterol content in Cav1KO MVBs induces changes in exosome membrane organization in these cells, we isolated cold- detergent-resistant microdomains (exosome DRMs) from the exosomes produced by Cav1WT cells (control or U18666A-treated) and Cav1KO cells. Flotillin-1, a well-known marker of exosomes and liquid ordered domains, showed a wider distribution in Cav1KO cells (fractions 1 to 8) or U18666A-treated Cav1WT cells (fractions 1 to 6); in contrast, flotillin accumulation in control Cav1WT-derived exosomes was restricted to fractions 1 to 3 (Fig. 2g), reflecting the increased high-ordered membrane content of Cav1KO-derived exosomes. Altered lipid composition as a major direct consequence of Cav1 deficiency in exosomes was further supported by mass spectrometry analysis of phosphatidic acid (PA), phosphatidylinositol (PI), and LBPA, which is essential for ILV budding from the MVB inner leaflet ^42^,^43^. Cav1WT-derived exosomes had significantly higher amounts than Cav1KO exosomes of these components (Supplementary fig. 2c).

These results strongly indicate that Cav1 plays a key role in the modulation of exosome biogenesis by acting as a “cholesterol rheostat” in MVBs, thus conferring plasticity to this compartment and enabling the segregation of different exosome populations.

### Cav1 promotes the sorting of specific ECM cargoes into exosomes

To determine whether Cav1-dependent heterogeneity in exosome size influences protein cargo composition, we analyzed proteins from Cav1WT and Cav1KO fibroblasts and their corresponding exosome preparations by isobaric labelling-based quantitative proteomics. This analysis identified a subproteome that is equally abundant in Cav1WT and Cav1KO whole-cell lysates but whose components are differentially sorted to exosomes depending on Cav1 genotype (Fig. 3a left panel). At a change threshold of |Zq| > 1.5, 152 proteins from this subproteome were more abundant and 163 proteins were less abundant in Cav1KO exosomes than in Cav1WT exosomes. Interaction network and functional annotation enrichment analysis revealed that Cav1WT-derived exosomes were significantly enriched in ECM components (including TnC, fibronectin, nidogen, emilin, EDIL3 and heparan sulfate proteoglycans) and endosomal/endomembrane system components (Fig. 3a right upper network, Fig3b, Supplementary Fig. 2d and Supplementary Table 1). Conversely, STRING network analysis revealed marked enrichment of Cav1KO exosomes for DNA/RNA binding and chaperone proteins such as histones (Fig. 3a right lower network, Fig 3b, Supplementary Fig. 2d and Supplementary Table 1). Consistent with our data supporting an increased cholesterol content in Cav1KO MVBs, various annexin family members were substantially enriched in Cav1KO exosomes, despite the slightly decreased abundance of these proteins in Cav1KO whole-cell lysates (Fig. 3c). Annexins are Ca2+-dependent proteins that bind cholesterol-rich membranes, providing lipid-clustering activity. Annexin enrichment in Cav1KO exosomes could may be a mechanism to sequester excess cholesterol in these exosomes.

**Figure 3.**
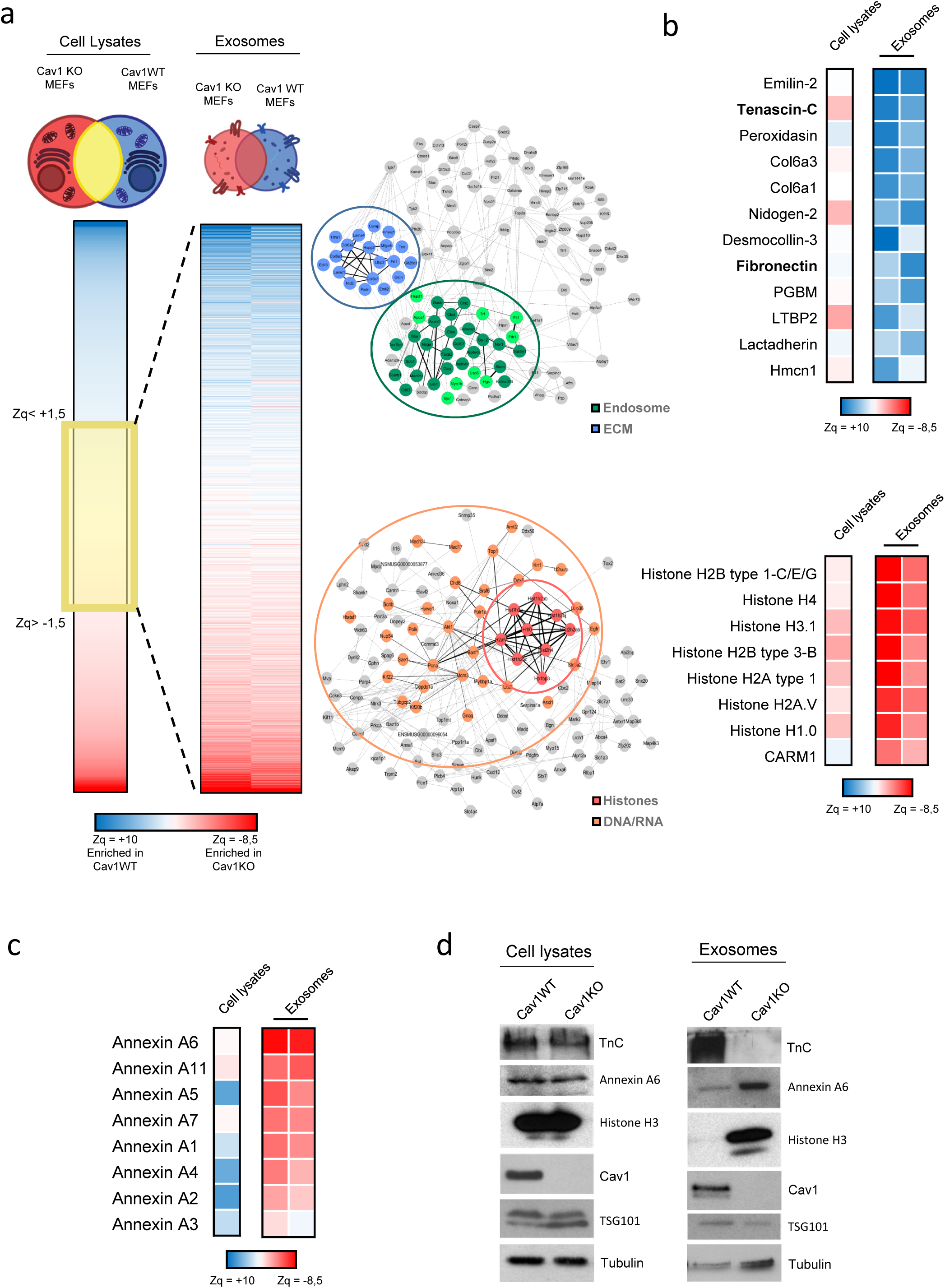
Cav1 specifies protein cargo into exosomes (a) Representation of hierarchical analysis of proteins equally expressed in the 2 parental cell populations (cell lysates, shaded in yellow) and differentially expressed in exosomes. *Zq* is the base 2 logarithm of protein ratios expressed in units of standard deviation, according to the WSPP model^69^,^70^. Blue indicates proteins upregulated in Cav1WT-MEF-derived exosomes, whereas red indicates proteins upregulated in Cav1^-/-^-MEF-derived exosomes. The graphics on the right show STRING network analysis. *Top.* Interactions among identified proteins upregulated in Cav1WT-MEF-derived exosomes. Highlighted groups are related to extracellular matrix components (blue) and extracellular vesicle/exosome biogenesis and lysosome components (green). *Bottom*. Interactions among identified proteins upregulated in Cav1KO-MEF-derived exosomes. Highlighted groups are related to histones (red) and DNA/RNA conformation and binding (orange). (b) Clustered heatmap of extracellular proteins with significant changes in extracellular matrix protein abundance between exosomes produced by Cav1WT and Cav1KO fibroblasts (top) or in histone related proteins (bottom). (c) Heatmap comparing annexin isoform enrichment in exosomes produced by Cav1KO and Cav1WT fibroblasts; colour code as in (a). (d) Western blot analysis confirming the protein expression changes identified by quantitative proteomics in Cav1WT and Cav1KO fibroblast lysates and derived exosomes.

The differential exosome sorting of selected gene products was validated by conventional western blot analysis (Fig. 3d). A particularly marked difference was observed for TnC (*Zq*=3.02), an extracellular matrix protein involved in development, stem cell niche formation, and aggressive tumour metastasis^44^-^46^. TnC was prominently represented in Cav1WT-derived exosomes but virtually absent from Cav1KO exosomes (Supplementary Table 1). Consistent with these observations, TnC colocalized with Cav1 within MVBs in fibroblasts and U251 cells (a TnC-secreting glioblastoma cell line^47^) (Supplementary Fig. 2e). These results strongly support the view that Cav1 modulates exosome biogenesis and the sorting into exosomes of specific cargo proteins, especially ECM components. The data also suggest that Cav1 is a prominent regulator of exosome-mediated intercellular communication, with as-yet uncharacterized downstream biological functions.

### Exosome biogenesis/secretion modulates ECM deposition

We next examined whether exosome-delivered ECM components can initiate *de novo* ECM deposition. Fluorescent immunolabelling and confocal microscopy revealed that Cav1KO fibroblast cultures deposit significantly less TnC fibre matrix in extracellular spaces than wild type cells, a result recapitulated by lentivirus-mediated knockdown in Cav1WT fibroblasts (Cav1KD) (Fig. 4a). TnC fibre deposition was also reduced by treatment of Cav1WT cells with the cholesterol trafficiking inhibitor U18666A (Fig. 4b and 4c and Supplementary Fig. 3b). Reduced TnC deposition associated with Cav1 downregulation or genetic deletion was concomittant with intracelular TnC accumulation; in contrast, U18666A-treated wild-type cells showed less evident intracellular TnC retention, probably reflecting a much higher rate of TnC degradation in this condition (Fig. 4b). Importantly, TnC was significantly less abundant in exosomes derived from Cav1KO and Cav1KD MEFs or U18666A-treated Cav1WT MEFs (Fig. 4d and Supplementary Fig. 2f), supporting a link between exosomes and TnC matrix deposition. To assess the contribution of exosomes to extracellular TnC fibre deposition, we treated Cav1WT fibroblasts with one of two available exosome secretion inhibitors: dimethylamiloride (dMA), an H+/Na+ and Na+/Ca2+ channel inhibitor, and GW4869, a neutral sphingomyelinase-2 (nSMAse2) inhibitor^36^,^48^. Net exosome release by Cav1WT fibroblasts was reduced by nSMAse2 blockade, and to a lesser extent by H+/Na+ and Na+/Ca2+ channel blockade (Supplementary Fig. 3a), correlating with an impaired ability to deposit extracellular TnC fibres (Fig. 4e). Both treatments also provoked the accumulation of dense intracellular TnC aggregates (Supplementary Fig. 3c). Importantly, the deposition of fibronectin (FN), another key ECM component underrepresented in Cav1KO exosomes (Fig. 3b and Supplementary table 1), was also reduced by exposure to exosome secretion inhibitors, with GW4869 having a much stronger effect (Fig. 4f). The general impact of exosome secretion on ECM deposition was also confirmed in primary cancer-associated fibroblasts (CAFs), a tumour stroma population with a key role in cancer progression. Exosome release by CAFs was largely inhibited by GW4869, whereas dmA had virtually no effect (Supplementary Fig. 4a). Accordingly, TnC and FN deposition by CAFs was specifically affected by nSMAse2 blockade, whereas dMA had no effect (Supplementary Fig. 4b and 4c). Confocal analysis and quantitative ER stress assays showed that neither GW4869 nor dMA affected the function or structure of ER-Golgi compartments in MEFs or CAFs (Supplementary Fig. 3d and 3e and Supplementary Fig. 4d and 4e), ruling out compromised ER-Golgi secretory function as a possible cause of the defective deposition of ECM components.

**Figure 4.**
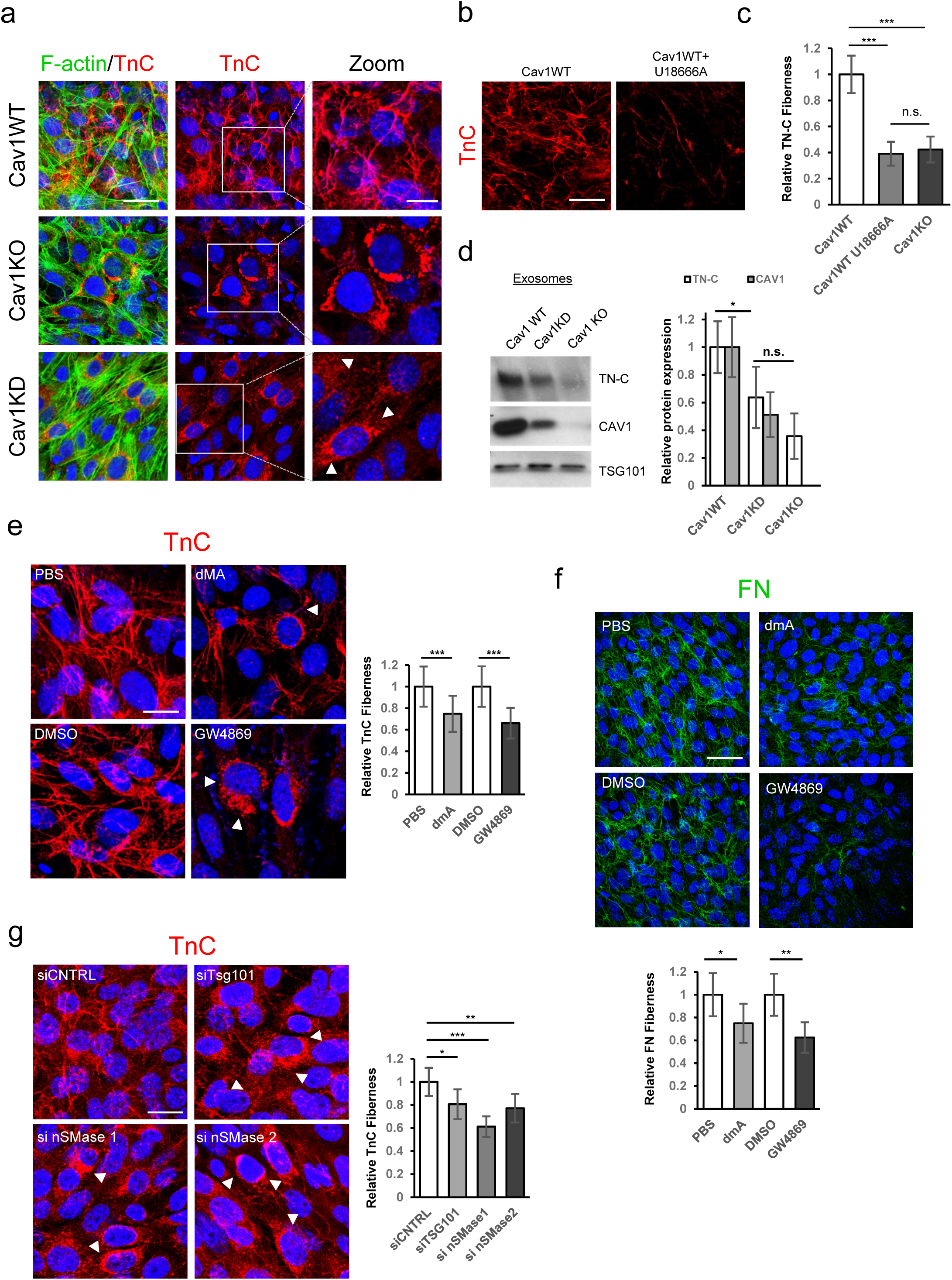
Exosome-mediated extracellular matrix deposition is caveolin-1 dependent Z-stack projection of confocal microscopy images showing TnC (red) in matrix deposited by Cav1WT, Cav1KD and Cav1KO MEFs. F-actin is shown in green (Scale bar, 40 µm). Zoomed views reveal an absence of TnC matrix deposition and a corresponding increased intracellular accumulation (white arrow) in Cav1KD and Cav1KO MEFs (Scale bar, 10 µm). TnC deposition of Cav1WT MEFs treated with U18666A determined by Z-stack projection confocal microscopy (Scale bar, 40 µm). (c) The chart shows extracellular TnC deposition in fibres produced by Cav1WT, Cav1KO and U18666A-treated Cav1WT MEFs, measured with in-house software (mean±s.d; n=8). (d) Western blot analysis of TnC downregulation in exosomes upon Cav1 depletion. TnC expression in exosomes secreted by Cav1-knockdown MEFs is compared with expression in exosomes derived from Cav1WT and Cav1KO counterparts. The chart shows TnC and Cav1 levels in each case (mean±s.e.m; n=6). (e) Representative immunofluorescence images showing the effect of 5-day exposure to the exosome inhibitors dimethylamyloride (dMA, 75 nM) and GW4869 (10 μM) on TnC matrix deposition (red) by Cav1WT MEFs (Scale bar, 20 µm). The chart to the right shows extracellular TnC deposition in fibres measured with in-house software (mean±s.e.m, n=12). (f) Representative Z-stack projection immunofluorescence confocal microscopy images showing the effect of 5-day exposure to dMA (75 nM) and GW4869 (10 μM) on FN matrix deposition (green) by Cav1WT MEFs (Scale bar, 50 µm). The chart to the right shows extracellular FN deposition in fibres measured with in-house software. Error bars are mean±s.e.m, n=5. (g) Representative Z-stack projection immunofluorescence confocal microscopy images showing the effect of siRNAs targeting Tsg101 and neutral sphingomyelinases 1 and 2 (nSMase1 and 2) on MEF TnC matrix deposition (red) (Scale bar, 20 µm). The chart to the right shows extracellular TnC deposition in fibres measured as in c (mean±s.e.m; n=6).

Because ESCRT machinery-regulated trafficking and ceramide biogenesis pathways are thought to generate exosomes with different cargoes^9^, we analyzed the effect on TnC matrix deposition of transient knockdown of Tsg101 (a core regulator of the ESCRT-dependent pathway) or of nSMAse1 and 2 (key regulators of ceramide metabolism) (Supplementary Fig. 3f and 3g). Consistent with the results obtained with small compound inhibitors (Fig.4e), knockdown of either pathway impaired TnC deposition; however, the phenotype was stronger upon ceramide synthesis disruption (Fig. 4g and Supplementary Fig. 3h). Similarly, the same knockdown conditions significantly reduced FN deposition (Supplementary Fig. 3i). These data support a role of exosomes as a general ECM deposition mechanism, with ceramide-dependent exosome biogenesis as a major limiting step in this process.

### TnC pools incorporated into exosomes have an intracellular origin

We next investigated the mechanisms underlying TnC incorporation into exosomes. To rule out an external orgin of exosome-incorporated TnC through endocytic trafficking, we developed an assay based on cell-derived matrices (CDMs). CDMs were generated by decellularization of ECMs synthesized *in vitro* by TnCWT or TnCKO MEFs. Immunostaining and microscopy analysis confirmed that TnCKO cells were specifically impaired for TnC deposition but not for deposition of other ECM components such as FN (Fig. 5a and Supplementary Fig. 4f). TnCKO or TnCWT cells were then seeded on each of these CDMs (Fig. 5b) and TnC internalization was determined by western blot. No intracellular TnC was detected inside TnCKO fibroblasts grown on TnC-rich CDMs, thus excluding endocytosis as an active TnC pool for exosome cargo sorting.

**Figure 5.**
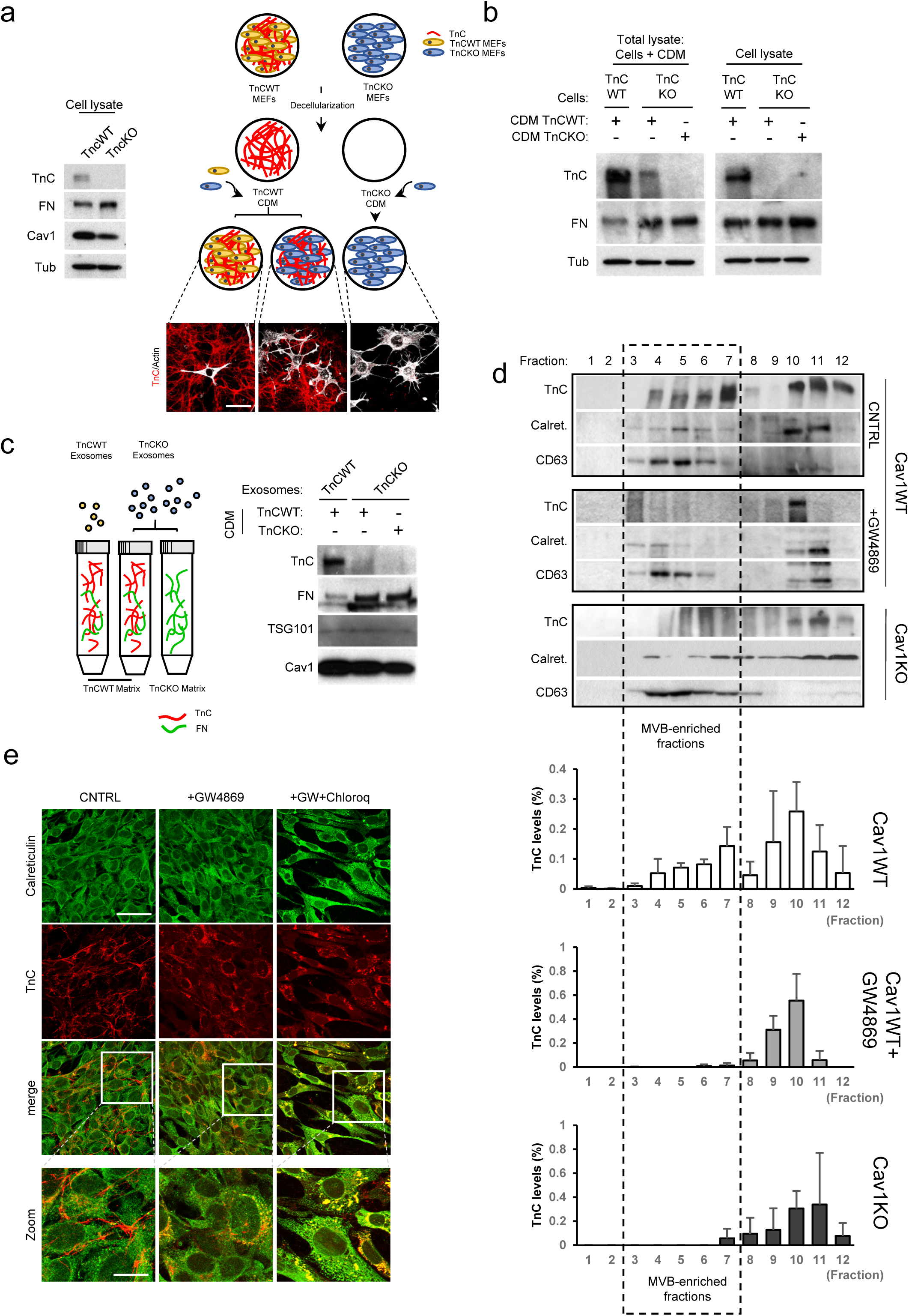
Exosomal incorporation of TnC occurs intracellularly via the ER-MVB route. (a) Western blot analysis of TnC and FN expression in lysates from TnCWT and TnCKO fibroblasts (left panels). Cell-derived matrices (CDMs) were generated from TnCWT and TnCKO cells as described in materials and methods, and TnCWT and TnCKO cells were plated on these CDMs as depicted. CDMs were labelled with TnC (in red) and the plated cells stained with F-actin (in grey) (Scale bar, 50 µm). (b) Western blot analysis of TnC and FN in total lysates (CDM+plated cells) and CDM-plated cells alone. Tubulin was used a loading control. (c) Interaction assay of purified exosomes with TnC rich or TnCKO CDMs. The scheme shows the assay protocol. The western blot shows analysis of TnC and FN binding to exosomes. (d) Subcellular fractionation analysis of intracellular TnC distribution in untreated Cav1WT fibroblasts, Cav1WT cells treated with GW4869, and Cav1KO cells. Western blot analysis was used to detect the MVB marker CD63 and the ER marker calreticulin. The boxed area denotes MVB-enriched fractions. The plots show the TnC present in MVB-enriched fractions in each condition (mean±s.d, n=3). (e) Colocalization analysis of TnC (red) and calreticulin (in green) in Cav1WT MEFs treated with GW4869 alone or in combination with the lysosomal inhitor chloroquine (Scale bar, 50 µm; zoomed views scale bar, 20 µm).

This finding left open the possibility that TnC detected in purified exosome fractions might originate in the passive, nonspecific binding of extant extracellular TnC fibres to the surface of secreted exosomes. To exclude this, purified exosomes from TnCKO cells were incubated with TnC-rich matrix produced by TnCWT cells (see scheme in Fig. 5c). After a second round of exosome purification, western blot analysis failed to detect TnC protein in TnCKO exosomes even after their incubation with TnC-rich matrix (Fig. 5c).

To further characterize TnC supply to exosomes, we conducted sucrose gradient ultracentrifugation analysis to separate MVBs from the endoplasmic reticulum (ER) and Golgi compartments (which fractionate together). In untreated Cav1WT cells, TnC co-sedimented with the MVB marker CD63 and the ER marker calreticulin (Fig. 5d); however, upon treatment with the exosome release inhibitor GW4869, TnC was largely excluded from CD63-positive fractions, a distribution shift recapitulated in Cav1KO-derived cell homogenates (Fig. 5d). Moreover, immunofluorescence analysis again revealed a major fraction of cellular TnC colocalizing with the ER marker in Cav1KO cells or GW4869-treated Cav1WT cells (Fig. 5e and Supplementary Figure 4g). Interestingly, this colocalization was significantly enhaced in the presence of the lysosomal inhibitor chloroquine. These results suggest that defective transfer of TnC from the ER to MVBs results in retention of TnC in the ER and its subsequent targeting for degradation, supporting the idea that lysosomal degradation is a mechanism of TnC removal when its sorting into exosomes is impaired. Degradation rate differences could explain the cell-cell variability in TnC accumulation in Cav1KO cells and in wild type cells treated with exosome inhibitor drugs and why severe disruption of exosome biogenesis and secretion does not entail a robust engagement of the unfolded protein response.

Together, these observations strongly support the view that the presence of TnC in exosomes reflects true regulated intracellular sorting of TnC into MVBs, likely through a mechanism involving ER-MVB contact sites, and that this exosomal TnC pool is sourced in the cell from *de novo* translation and not from endocytosed material.

### Cav1-dependent exosome biogenesis directy drives ECM deposition and subsequent increase in tumour-cell invasiveness

We next compared the ability of purified exosomes derived from either Cav1WT or Cav1KO MEFs to deposit TnC on 2D cultures of the non-invasive breast cancer tumour cell line MDA- MB468, which naturally expresses negligible levels of Cav1 and TnC^49^. After 24h, tumour cells internalized pre-labelled Cav1WT and Cav1KO exosomes with similar efficiency, as shown by similar accumulation of the general membrane dye PKH67 (Fig. 6a, in green). However, only Cav1WT-derived exosomes led to detectable deposition of TnC in areas close to the periphery (Fig. 6a, in red). In contrast, MDA-MB468 cultures exposed to Cav1KO exosomes showed baseline levels of TnC deposition comparable to those displayed by control cultures (exposed to vehicle). Exosomes thus have an intrinsic ability to deposit specific ECM material *in vitro* that requires Cav1-dependent regulation of exosome biogenesis, likely derived from the specification of cargo profile.

**Figure 6.**
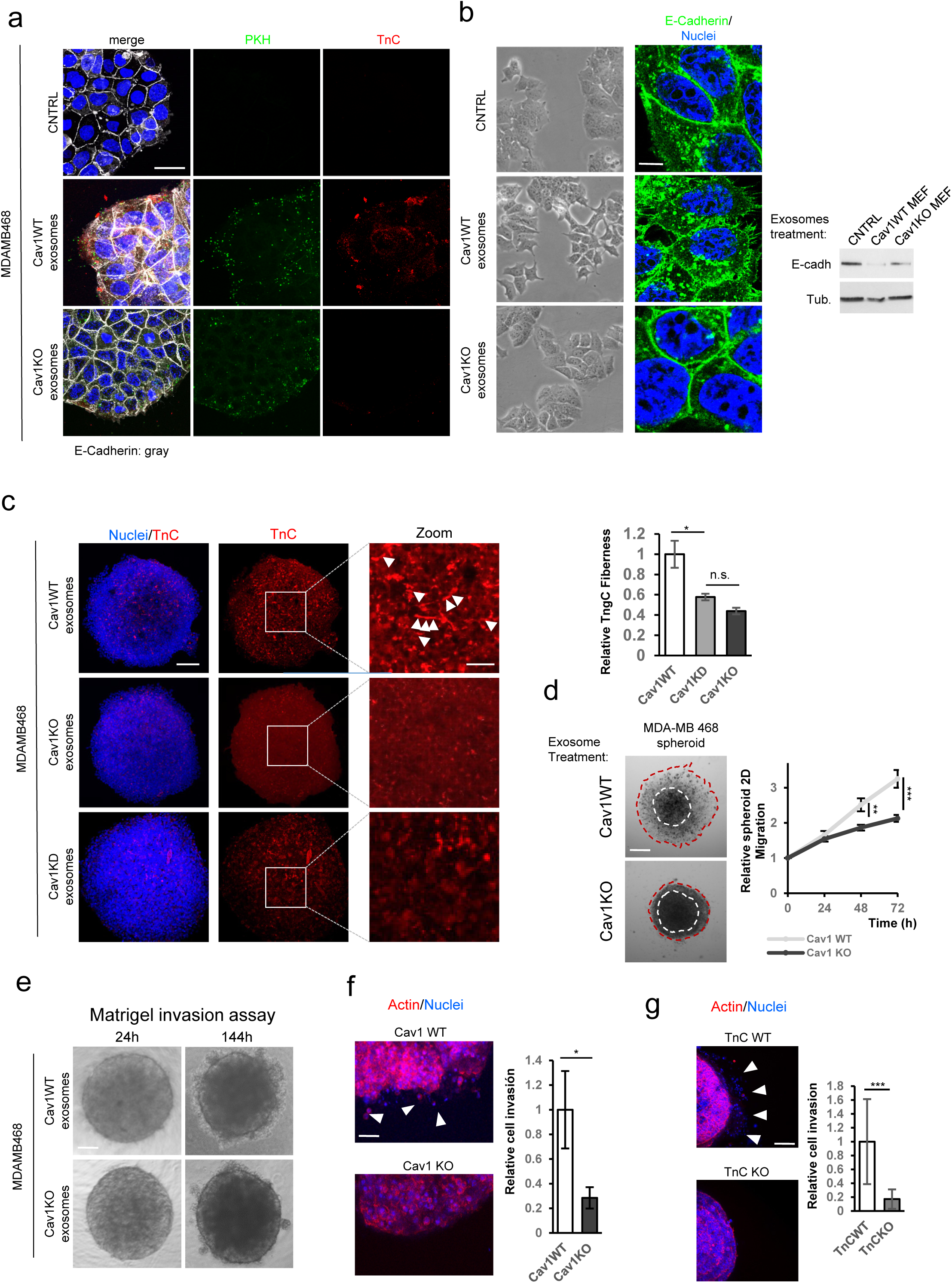
TnC deposition mediated by exosomes promotes tumour cell invasiveness TnC deposition by exosomes was determined in 2D culture conditions. MDA-MB468 breast tumour cells were incubated with PKH67-labelled exosomes derived from Cav1WT or Cav1KO MEFs or with PBS (control). (a) Confocal microscopy analysis of exosome (green) uptake by MDA-MB468 cells and the deposition of exosome-delivered TnC (red) in areas surrounding the tumour cells. White arrowheads indicate TnC deposits. (Scale bar, 30 µm) (b) Phase-contrast and confocal microscopy analysis of morphological changes in MDA-MB468 tumour cells 36 h after exposure to fibroblast-derived Cav1WT and Cav1KO exosomes. Cells were stained for the EMT marker E-cadherin; nuclei were stained with Hoescht. Zoomed images show E-Cadherin redistribution from cell-cell contact sites to intracellular compartments (Scale bar 10 µm). E-Cadherin was determined by western blot. (c) Z-stack confocal microscopy analysis of TnC deposition (red) within 3D MDA-MB468 spheroids generated in the presence of Cav1WT, Cav1KO or Cav1KD exosomes. Nuclei are stained with Hoechst (blue) (Scale bar, 150 µm). Zoomed images show a clear distribution of TnC deposits (Scale bar, 50 µm). White arrows indicate the presence of TnC fibres. The chart to the right shows extracellular TnC deposition in fibres into the spheroids (mean±s.e.m, n=3). (d) Phase contrast microscopy of MDA-MB468 spheroid migration (Scale bar, 175 µm). Spheroids were generated in the presence of the indicated exosomes, transferred to 96-well plates, and maintained in culture for 72 h. The graph shows quantitative analysis of spheroid migration (mean±s.e.m; n=11). (e) Invasiveness of MDA-MB468 spheroids generated in the presence of fibroblast-derived Cav1WT and Cav1KO exosomes. Invasiveness was assessed in matrigel by phase-contrast microscopy (Scale bar, 150 µm). (f) and collagen type I gel by immunofluorescence Z-stack confocal microscopy. Arrows indicate areas of cell invasion. Quantification of invasiveness of MDA-MB468 spheroids into collagen is shown (mean±s.e.m; n=24 spheroids per condition) (Scale bar, 100 µm). (g) Invasiveness of MDA-MB468 spheroids generated in the presence of fibroblast-derived TnCWT and TnCKO exosomes determined by confocal microscopy. Arrows indicate areas of cell invasion. The chart shows quantification of MDA-MB468 spheroid invasiveness into collagen (mean±s.d; n=15 spheroids per condition; scale bar, 150 µm).

The deposition of TnC delivered by Cav1WT-derived exosomes induced important morphological changes in receptor MDA-MB468 tumour cells, eliciting cell-cell adhesion loss and internalization of the epithelial marker E-cadherin, associated with increased protrusiveness and emission of filopodia (Fig. 6b). A similar phenotype was produced by using CAF lines transduced with lentiviral vector encoding Cav1 shRNA (Supplmentary Figure. 5b). Consistent with these EMT-like changes, MDA-MB468 cells concomitantly acquired an increased ability to migrate, as determined in wound healing studies and transwell cell migration assays (Supplementary Fig. 5c and 5d). In contrast, exposure of MDA-MB468 cells to purified exosomes derived from Cav1KO fibroblasts induced no significant morphological changes (Fig. 6a and 6b). Interestingly, exosomes purified from MDA-MB231 tumour cells, which contain Cav1 but not TnC, also failed to induce an increase in MDA-MB468-cell protrusiveness and motility (Supplementary Fig. 5a and Supplementary Table 2), strongly indicating that exosomal TnC/ECM cargo, and not Cav1, is the direct driver of the phenotypic alterations.

The *in vivo* setting is more closely modelled by 3D culture methodologies^50^. We thus studied the ability of exosomes to deposit ECM on MDA-MB468 spheroids. Spheroids are scaffold-free 3D assemblies in which cells self-organize ECM and engage in intercellular communication; spheroids have been validated as superior *in vitro* experimental proxies of live tissues for many applications^50^,^51^. Consistent with the 2D culture experiments, Cav1WT- derived TnC-containing exosomes, but not exosomes derived from Cav1KO or Cav1KD fibroblasts (i.e. TnC-depleted), efficiently deposited fibrillar TnC structures within cell spheroids (Fig. 6c). 3D-reconstruction of confocal optical sections across the whole spheroid volume demonstrated that TnC fibrils were preferentially located in the interstitial space, enveloped by the peripheral actin of the cell membranes (Supplementary Fig. 5e). These experiments further support the view that exosomes from Cav1WT MEFs carry ECM components and are able to transfer these components to tumour cells for the deposition of new ECM.

To characterize Cav1-dependent, exosome-induced phenotypic changes in tumour cells, we monitored the boundary of MDA-MB468 spheroids over 72 h. Spheroids generated in the presence of Cav1WT-derived exosomes were seeded onto a plate and cells were allowed to migrate away from the spheroid. Cav1WT MEF-derived exosomes promoted significantly more migration than exosomes derived from Cav1KO MEFs (Fig. 6d). To confirm the role of TnC in this process, we tested the effect of exosomes derived from TnCKO MEFs (Supplementary Fig. 5f and 5g). TnCKO exosomes were poor stimulators of cell migration, supporting the view that the ability of Cav1WT MEF-derived exosomes to induce phenotypic changes in target tumour cells derives largely from Cav1-dependent sorting of specific ECM cargoes to exosomes.

We next examined whether exosome-induced tumour-cell protrusiveness and migration was associated with increased invasiveness through different substrates. Compared with exosomes derived from other sources, Cav1KO exosomes consistently exhibited a blunted ability to induce invasiveness in Matrigel invasion assays (Fig. 6e) or 3D collagen matrix invasion assays (Fig. 6f), similar to the reduced invasion promoted by exosomes derived from TnCKO cells (Fig. 6g)). These results further support the relevance of Cav1- and TnC-dependent exosome-induced ECM deposition and phenotype alteration for tumour progression.

### Cav1WT MEF-derived exosomes create TnC-rich niches *in vivo*

We next evaluated whether exosomes deposit TnC *in vivo*, surviving vascular dissemination over large (inter-organ) distances. We fluorescently labelled the membranes of exosomes purified from Cav1WT or Cav1KO fibroblasts and injected into the caudal veins of TnCKO mice every day for 1 week (Fig. 7a). Histopathological analysis revealed specific exosome accumulation in liver, and to a lesser extent, in lung (Fig. 7b and Supplementary Fig. 5h). However, associated TnC deposition was only consistently detectable in liver and lung of mice injected with Cav1WT-derived exosomes, contrasting the residual TnC staining in the organs of animals injected with Cav1KO exosomes or vehicle (PBS), despite Cav1KO derived-exosomes being able to reach to this organ with comparable efficiency. Intriguingly, TnC from Cav1WT exosomes tended to accumulate preferentially in liver sinusoids (Fig. 7b), whereas in the lung TnC deposition was concentrated in areas surrounding mesenchymal cells (Supplementary Fig. 5h). Moreover, Cav1WT-derived exosomes showed a moderate preference to accumulate in liver rather than lungs, whereas Cav1KO exosomes showed no such preference, despite efficient uptake across organs. These organ-specific peculiarities might indicate a Cav1-dependent role for exosomes in targeting a specific cell subpopulation for niche formation.

**Figure 7.**
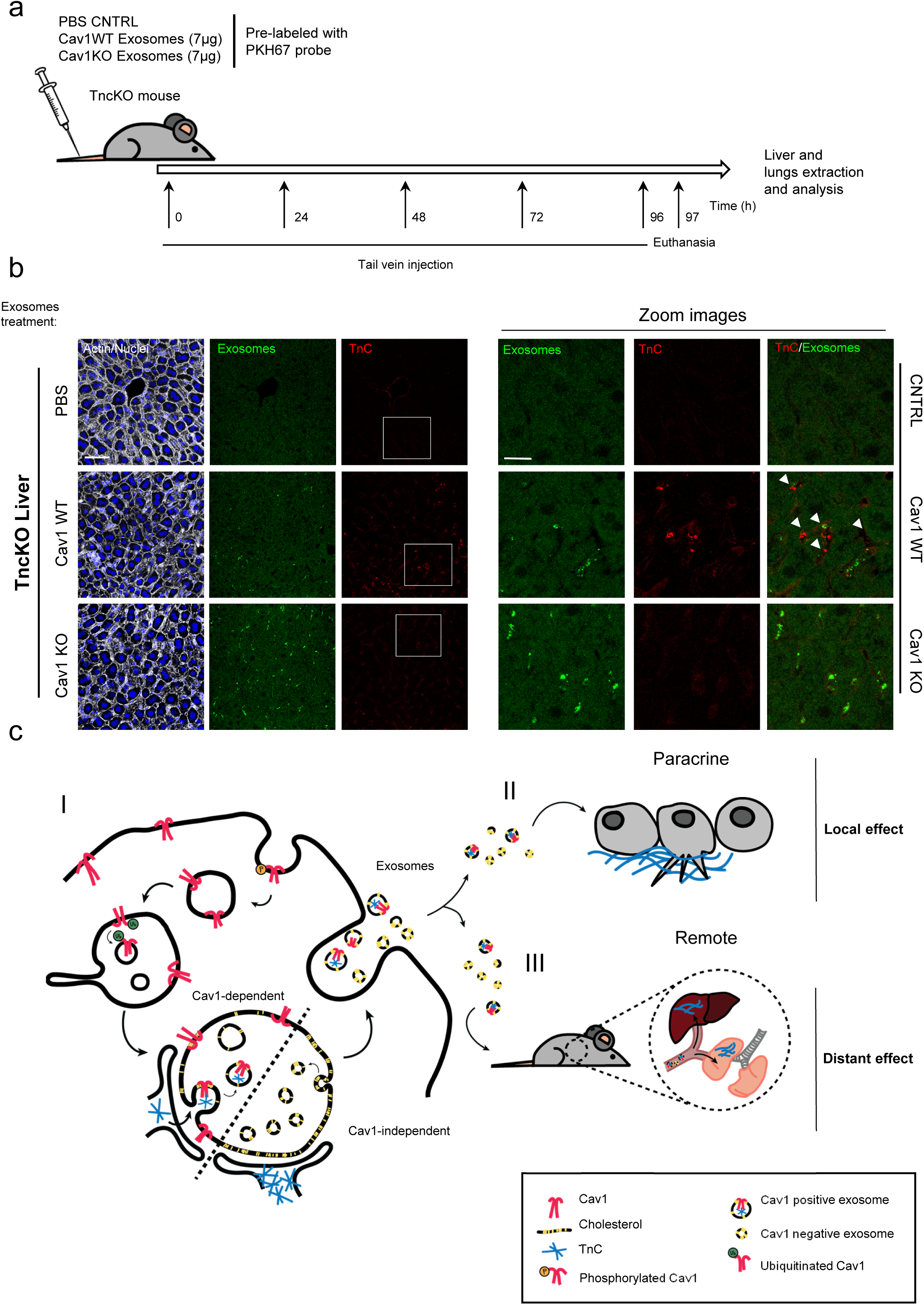
Cav1-loaded fibroblast-derived exosomes deposit TnC *in vivo* in a tissue specific manner (a) Treatment protocol. (b) Fluorescence microscopy of liver sections from mice injected in the tail vein with Cav1WT or Cav1KO fibroblast-derived exosomes or with PBS (Scale bar, 60 µm; zoomed views, scale bar, 30 µm). Exosome foci are green. White arrows indicate regions of TnC deposition. All images are representative of nine random fields from two independent experiments. (c) Proposed role of Cav1 in exosome-mediated ECM deposition. (I) Endocytosed Cav1 enters the MVB compartment, where it promotes exosome heterogeneity in size and composition, favouring the entry of specific ECM components. (II) Cav1WT fibroblast-derived exosomes stimulate protrusive activity and motility of breast cancer cells by nucleating local extracellular matrix (TnC) deposition in areas surrounding the tumour cells. (III) Cav1WT fibroblast-derived exosomes also generate extracellular matrix-rich deposits at long distances from their source *in vivo*, generating sites of possible future metastasis.

Our observations thus support a key role for Cav1 in exosome-dependent ECM deposition *in vivo*. TnC is a positive regulator of metastasis through the assembly of favourable distant microenvironments^52^, suggesting that Cav1 might specifically modulate cancer outcome through exosome-dependent stroma-tumour communication.

## DISCUSSION

Exosomes have recently attracted attention due to their potential to provide a means by which tumour cells and associated stroma not only communicate locally, but also influence and transform distant locations or ‘niches’ and even modulate organismal responses systemically^12^-^14^. Despite the ubiquity of exosomes, the mechanisms underlying exosome biogenesis, cargo sorting, and function remain elusive. Here, we demonstrate the role of Cav1 as a modulator of exosome biogenesis through the regulation of cholesterol content in MVBs. Cav1-dependent cholesterol modulation in this compartment permits the generation of exosomes with a higher heterogeneity, specifying exosome cargo subsets (mainly specific ECM proteins) and subsequent functional impacts. Our data demonstrate that the presence of Cav1 within exosomes is favoured by its endocytosis through specific phosphorylation at tyrosine 14. This phosphorylation step promotes Cav1 targeting to the MVB outer membrane, where a second mechanism is activated that requires the presence of ubiquitination-targeting N-terminal domain lysine residues; this second mechanism sorts Cav1 into ILVs for exosome generation. By virtue of its cholesterol-binding ability, Cav1 would act as a “cholesterol rheostat” in MVBs, thus increasing the plasticity of this compartment and enabling the segregation of different exosome populations, a phenomenon that as we have demonstrated can be perturbed by pharmacological treatments that directly affect intracellular cholesterol trafficking. The importance of cholesterol regulation in this compartment could also determine how exosome protein cargo is specified through modulation of contact between the ER and MVBs. (Fig. 7c, enlarged graph).

Among the exosome-targeted ECM proteins identified, TnC is a key component involved in promoting exosome-dependent tumour invasion. TnC matrix deposition mediated by Cav1- expressing fibroblasts requires intact exosome biogenesis and secretion, as demonstrated by the effect of exosome inhibitors and siRNAs targeting exosome biogenesis-machinery components. Furthermore, our data support a model whereby the ER/Golgi compartment is a major direct supplier of TnC to these exosomes.

Exosomes secreted by Cav1WT fibroblasts have a dual effect, on the one hand stimulating protrusiveness and active migration of targeted tumour cells by inducing an EMT-like process, and on the other hand eliciting ECM deposition in areas surrounding the tumour. We demonstrate that this effect also takes place *in vivo* over large distances, suggesting that these mechanisms contribute to tumour-cell invasiveness and favour the establishment of distant metastatic foci by nucleating new ECM deposition (Fig. 7c).

Cav1 is trafficked through several endomembrane systems, including late endosome/MVBs and lysosomes, suggesting the likely existence of several as yet unidentified caveolae-independent Cav1 functions ^24^,^53^. Previous studies demonstrated that caveolae budding from the plasma membrane carry Cav1 to early endosomes, presumably allowing dynamic recycling and maintenance of these structures^54^. However, an alternative pathway has been described in which caveolae are disassembled and Cav1 is internalized and ubiquitinated, followed by accumulation at the internal membrane of late endosomes/MVBs^32^,^35^. Interestingly, Cav1 is also targeted to late endosomes by perturbations of lysosomal pH and changes in cholesterol content, with no changes in Cav1 degradation rate, raising the possibility of alternative fates for this Cav1 pool ^33^. Our data demonstrate that a significant pool of endocytosed Cav1 is sorted into MVBs en route to exosomes (Fig. 1), but we cannot rule out that other intracellular compartments such as the ER may also serve as a source of exosomal Cav1 (Fig. 1 and Supplementary Fig. 1).

Cav1 binds cholesterol with high affinity ^40^ and its ability to move between different compartments might contribute to the regulation of cholesterol fluxes and distribution within cells^55^. Our data demonstrate that Cav1 absence increases the level of free cholesterol in late endosomes/MVBs, suggesting that Cav1 is important for the modulation of membrane plasticity required for the generation of ILVs of varying size. ILV size heterogeneity would favour a specific and significant enrichment of extracellular matrix components in exosomes. (Fig. 2 and Supplementary Fig. 2 and Fig.3). Interestingly, among the identified Cav1- dependent exosomal cargoes, collagens are less abundant than other matrix proteins such as TnC, fibronectin, laminin, nidogen, and emilin. Collagen secretion requires specific assembly mechanisms operating through the COPII biosynthetic/secretory machinery^56^, and is thus unavailable to the exosome pathway. (Fig3, Fig. 4 and Supplementary Fig. 3). These pathways might be oppositely regulated by Cav1, since a negative correlation has been described between Cav1 expression and collagen deposition in chronologically-aged skin and during fibrosis^57^,^58^. Our results strongly suggest that a previously overlooked critical function of Cav1 is ECM secretion via exosomes. Supporting our observation that exosomes can act as carriers of some extracellular matrix proteins, a recent study demonstrated exosomal secretion of FN through its binding to integrin α5β1; however, the function and underlying mechanism of this sorting were not determined.^59^,^60^

In constrast to Cav1WT-derived exosomes, the homogeneous exosome population produced by Cav1null cells has a strong tendency to aggregate (data not shown), which could be directly linked to the ability of annexins, significantly enriched in Cav1null exosomes, to favour cholerestol-enriched membrane aggregation^61^. Intriguingly, in the absence of Cav1, the main exosome protein components are related to DNA/RNA binding and nuclear functions, cargoes that may be favoured in conditions of metabolic stress (Fig. 3b).

Our data demonstrate the importance of TnC secretion via exosomes and point to incorporation of TnC into ILVs through its sorting from the biosynthetic route to the MVB compartment, excluding the possibility of an extracellular origin through its internalization into the cell or by direct binding to exosomes once they are released (Figure 5).

Recent work has revealed the presence of contact sites between MVBs and the ER^62^,^63^, but the frequency, dynamics and physical architecture of these contacts are poorly understood. Interestingly, our group has demonstrated the importance of Cav1 as a main component of a section of the smooth ER that forms close contacts with mitochondria^23^, suggesting that Cav1 may also modify ER-MVB contacts sites, thus modulating the incorporation of specific protein cargo into MVBs en route to exosomes.

The similar effects of TnCKO and Cav1KO exosomes (Fig. 6) suggest that the biological actions determined by secretion of Cav1WT-derived exosomes directly depend on exosome TnC content. Our results suggest that TnC has a dual effect of on tumour progression. Over short distances, fibroblast-derived TnC-rich exosomes are deposited in areas surrounding tumour cells. The presence of TnC at the tumour invasive front presumably induces an EMT-like process that drives tumour cell migration away from the bulk tumour mass, as evidenced in 2D and 3D assays (Fig. 6). Similar results were obtained by direct addition of recombinant soluble TnC^64^. Together with this local effect, our data provide evidence for exosome-mediated TnC deposition over large distances to sites in lung and liver. Recent findings demonstrate that the microenvironment required for the formation of secondary metastatic sites is primed before the arrival of cancer cells, forming a “landing dock” for future metastatic growth^65^. Establishment of a pre-metastatic niche is a multistep process involving the sequential activation of several cell types, as recently shown for metastasis of pancreatic cancer to liver^13^. We propose that this orchestrated cascade is complemented by alternative mechanisms that directly promote pre-metastatic niche formation. Our results show that caudal vein injection of exosomes derived from Cav1WT fibroblasts can promote direct TnC-matrix deposition in distant organs (Fig. 7).

In addition to the role of exosomes in tumour progression, altered extracellular vesicle secretion is also linked to several cholesterol-related diseases ^66^,^67^. This attracts particular attention in relation to atherosclerosis, where exosomes may be involved in monocyte and macrophage cholesterol metabolism, endothelial cell and platelet activation, and smooth muscle proliferation. Interestingly, Cav1 loss results in hypercholesterolemia but also protects against the development of aortic atheromas. Further investigation will be required to confirm whether modulation of exosomal Cav1 incorporation might provide an alternative way to modulate cholesterol levels as well as thre specific proatherogenic molecule cargo. The lysosomal storage disorder Niemann-Pick Type C (NPC) disease is also characterized by cholesterol accumulation in the endosomal-lysosomal system ^68^. Whether exosomes can serve as diagnostic and therapeutic tools for NPC patients remains to be determined. Although the specific connections between cholesterol homeostasis and exosomes remain unclear, Cav1 appears to be a key player in this interaction.

Previous studies demonstrated high levels of Cav1-expressing exosomes in the plasma of melanoma patients ^27^. We propose that relative levels of Cav1-expressing exosomes could be a valuable prognostic marker of metastasis and tumour malignancy as well as another cholesterol-related diseases. Our findings suggest new avenues for diagnosis, prevention, and therapeutic intervention based on specifically targeting Cav1-dependent roles in exosomal tumour-stroma communication.

## MATERIALS AND METHODS

### Antibodies and reagents

Monoclonal anti-caveolin-1 (#3267) and anti-Alix (#2171) antibodies were from Cell Signaling Technology. Anti-LBPA was from Echelon (Z-SLBPA). Anti-CD63 (ab1318), anti-flotillin (ab13349), anti-periostin (ab14041), anti-Calreticulin (ab2907), anti-PTRF (ab48824) and anti- TSG101 (ab83) were from Abcam. Anti-GM130 (610823), anti-py14-caveolin-1 (611339) and anti-Cadherin (610181) were from BD transduction. Anti-tubulin (T-9026), anti-Fibronectin (F3648) and anti-tenascin (Clone Mtn-12, T3413) were from Sigma. Anti-Tenascin was also from Millipore (AB19011). Anti-nSMAse1 (sc-377135) and anti-nSMAse2 (sc-166637) were from Santa Cruz. Alexa fluor 488, 546 and 647-conjugated phalloidin were from Life Technologies. GW4869 (567715) was from Calbiochem and dimethylamiloride (dMA) (A4562) was from Sigma. Matrigel (354230) was from BD and Collagen I from rat tail (354249) was from Corning. Filipin was from Sigma (F4767).

### Cell culture

Mouse embryonic fibroblasts derived from Cav1WT and Cav1KO littermate mice used throughout this study were kindly provided by Dr. Michael P. Lisanti. TnCWT and TnCKO MEFs were provided by our collaborator Dr. Gertraud Orend. PTRFKO cells were provided by Dr. Parton and were reconstituted with the retroviral expression vector pMIGR1. COS7 cells were from ATCC. MEFs and COS7 cells were cultured in Dulbecco´s modified Eagle medium (DMEM) supplemented with 10% fetal bovine serum (FBS), 100 U/ml penicillin and 100 µg/ml streptomycin at 37°C in a humidified 5% CO_2_-95% air incubator. The breast cancer associated fibroblast line CCD-1069S and the breast tumour cell line MDA-MB468 were cultured in DMEM/F-12 supplemented with 10% FBS, 100 U/ml penicillin and 100 µg/ml streptomycin. CCD-1069S fibroblasts stably expressing short hairpin RNA (shRNA) targeting Cav1 or a scramble siRNA were generated by lentiviral infection as reported in Goetz et al., 2011.

### siRNA silencing

siRNA silencing was performed using 20 nM siRNA and Lipofectamine RNAiMAX Reagent (Thermo Fisher Scientific) according to the manufacturer´s protocol, and cells were cultured for 2-3 days. siRNAs targeting mouse Tsg101, nSMAse1 and nSMAse2 were SMARTpool ON-TARGET plus reagents from Dharmacon (Tsg101 Cat. L-049922-01-0005; nSMAse1 or Smpd2 Cat. L-044206-01-0005; nSmase2 or Smpd3 Cat. L-059400-01-0005). The control siRNA was ON-TARGET plus non-targeting pool (Dharmacon Cat. D-001810-10-05). Cav1 was silenced with a siRNA from Dharmacon (seq. GAGCUUCCUGAUUGAGAUU) and an esiRNA from Sigma (EHU031251).

### Isolation and characterization of exosomes released by fibroblasts

Exosomes were isolated from cultured fibroblasts grown in exosome-free culture medium. To remove detached cells, conditioned medium was collected and centrifuged at 300 g for 10 min at 4°C. The supernatant was collected and centrifuged at 2000 g for 20 min at 4°C. The supernatant was then centrifuged at 10000 g at 4°C for 30 min to completely remove contaminating apoptotic bodies, microvesicles and cell debris. The clarified medium was then ultracentrifuged at 110,000 g at 4°C for 70 min to pellet exosomes. The supernatant was carefully removed, and crude exosome-containing pellets were washed in ice-cold PBS. After a second round of ultracentrifugation, the resulting exosome pellets were resuspended in the desired volume of PBS.

For further purification, exosomes were ultracentrifuged on a discontinuous sucrose gradient including sucrose concentrations of 0.25M, 0.5M, 0.8M, 1.16M, 1.3M and 2M in 20 mM HEPES, pH 7.4. Exosome samples were laid on the bottom of the gradient in the 2M sucrose fraction, followed by centrifugation at 200,000 g for 18 h. Eleven individual 1 ml gradient fractions were manually collected. Fractions were diluted in PBS and centrifuged at 110,000 g for 1 h at 4°C, and the resulting pellets were resuspended and analysed. Alternatively, an equal volume of cold acetone was added to each fraction and the proteins precipitated for 2 h at −20°C. Protein pellets were collected by centrifugation at 16,000 g for 10 min and air dried to eliminate acetone traces. The protein precipitates were monitored by western blot for the expression of the exosomal markers Alix and Tsg101.

In all experiments, the exosomes used corresponded to the total exosome pellets resulting from the serial centrifugation steps; usually 10^6^ exosome particles were added per cell. Exosome uptake was performed in exosome-free medium.

Exosome concentrations and size distributions were determined by Nanoparticle Tracking Analysis (Nanosight).

### Western blot analysis

Western blot samples were prepared with conventional 5X Laemmli sample buffer brought to 1X when mixed with the sample. The 5X Laemmli sample buffer is composed of 60 mM Tris-HCl pH 6.8, 2% SDS, 10% glycerol, 5% β-mercaptoethanol, and 0.01% bromophenol blue.

Cell or exosome proteins were resolved by SDS-PAGE and transferred to nitrocellulose membranes. Blots were incubated for 2 h in PBS containing 5% skimmed milk and overnight with primary antibodies (typically diluted 1:1000). After incubation (2 h) with horseradish peroxidase-conjugated goat anti-rabbit or goat anti-mouse antibody, signal on washed blots was detected by enhanced chemiluminescence (GE Healthcare). Band intensities were quantified with Image J software.

### Intracellular lysate analysis

To prepare lysates containing only proteins from the cell interior, plated cells were treated with trypsin to eliminate proteins in the extracellular space. Detached cells were washed with PBS and resuspended in loading buffer for western blot analysis. For total lysates comprising cell and ECM proteins, cells were scraped directly from the plate in loading buffer.

### PKH67 labelling of exosomes

Purified exosomes were labelled with the green fluorescent dye PKH67 (Sigma MIDI67) according to the manufacturer´s protocol. After staining, unbound PKH67 was removed with columns from Thermo Fisher (4484449).

### Electron microscopy

Exosomes were visualized by transmission electron microscopy (TEM) according to the method of Thery et al.^71^. The exosome suspension was fixed in 2% paraformaldehyde and transferred to formvar/carbon coated electron microscopy grids. After 20 minutes, grids were placed sample-face down for 2 minutes in a 100uL drop of PBS on a sheet of parafilm. Grids were then transferred to 1% glutaraldehyde for 5 minutes and then washed with distilled water. Samples were contrasted in 2% uranyl acetate and examined by TEM.

### Spheroid formation assay

After trypsinization, MDA-MB468 cells were resuspended in exosome-free DMEM, counted and adjusted to the desired concentration (usually 5×10^3^ cells/ml). A 20 µL final volume was prepared with 5X methyl cellulose prepared in exosome-free medium, the MDA-MB468 cell suspension, and the desired number of exosomes (usually 1X10^6^ particles). The mixture was carefully deposited in a drop on the inner surface of a 100 mm dish lid. The plate lid was placed on a plate containing 10 ml PBS to humidify the culture chamber. Under gravity, cells aggregated at the bottom of the hanging drop. After 24 hours, the resulting cell aggregates were lifted with a pipette. For 2D spheroid migration assays, the aggregates were seeded on a 96 well plate to monitor cell migration from the spheroid by phase-contrast time-lapse video microscopy. Alternatively, aggregates were seeded on collagen or matrigel gels as described below. In all assays, no additional exosomes were added to the medium apart from the exosomes used during spheroid generation.

### Collagen embedding

High-density rat-tail type I collagen (Corning) was diluted to 2 mg/ml. This solution self-polymerizes to form a gel after 20 to 30 min at 37 °C. Tumour cell spheroids were transferred to 25 µL unpolymerized collagen in the wells of a μ slide angiogenesis plate (IBIDI). After 30 min, 40 µL medium was added over the top of the gel. Medium was replaced every 48 h. Spheroids were imaged every 24 hours for the duration of the experiment. Spheroid diameter was measured from captured images with ImageJ.

For matrigel embedding assays, spheroids were transferred to a 25 µL 1:1 mix of matrigel and medium.

### Wound closure assay

For wound healing assay, confluent monolayers were scraped with a 0.1-2 μl pipette tip in the presence of mitomycin C, and wound closure was monitored by capturing images at 10-min intervals.

### Cell-derived matrix generation

CDM generation was as described in Goetz et al., 2011.

### Proteomic analysis

Lysates and exosomes derived from WT and Cav1^-/-^ MEFs were digested using the filter aided sample preparation (FASP) protocol^72^. Briefly, samples were dissolved in 50 mM Tris-HCl pH 8.5, 4% SDS and 50 mM DTT, boiled for 10 min, and centrifuged. Protein concentration in the supernatant was measured with the Direct Detect^®^ Spectrometer (Millipore). About 100 μg of protein were diluted in 8 M urea in 0.1 M Tris-HCl pH 8.5 (UA) and loaded onto 30 kDa centrifugal filter devices (FASP Protein Digestion Kit). The denaturation buffer was removed by washing three times with UA. Proteins were then alkylated by incubation in 50 mM iodoacetamide in UA for 20 min in the dark, and excess alkylation reagents were eliminated by washing three times with UA and three additional times with 50 mM ammonium bicarbonate. Proteins were digested overnight at 37°C with modified trypsin (Promega) in 50 mM ammonium bicarbonate at a 50:1 protein:trypsin (w/w) ratio. The resulting peptides were eluted by centrifugation with 50 mM ammonium bicarbonate (twice) and 0.5M sodium chloride. Trifluoroacetic acid (TFA) was added to a final concentration of 1% and the peptides were finally desalted onto C18 Oasis-HLB cartridges and dried-down for further analysis.

#### Multiplexed isobaric labelling

For the quantitative analysis, tryptic peptides were dissolved in triethylammonium bicarbonate (TEAB) buffer, and the peptide concentration was determined by measuring amide bonds with the Direct Detect system. Equal amounts of each peptide sample were labelled using the 4-plex iTRAQ Reagents Multiplex Kit (Applied Biosystems). Briefly, each peptide solution was independently labelled at room temperature for 1 h with one iTRAQ reagent vial (mass tag 114, 115, 116 or 117) previously reconstituted with ethanol. After incubation at room temperature for 1 h, the reaction was stopped with diluted TFA and peptides were combined. Samples were concentrated in a Speed Vac, desalted onto C18 Oasis-HLB cartridges, and dried-down for mass spectrometry analysis.

#### Mass spectrometry

Digested peptides were loaded into the LC-MS/MS system for on-line desalting onto C18 cartridges and analysed by LC-MS/MS using a C-18 reversed phase nano-column (75 µm I.D. × 25 cm, 2 µm particle size, Acclaim PepMap RSLC, 100 C18; Thermo Fisher Scientific) in a continuous acetonitrile gradient consisting of 0-30% B for 240 min and 50-90% B for 3 min (A= 0.5% formic acid; B=90% acetonitrile, 0.5% formic acid). A flow rate of 200 nl/min was used to elute peptides from the RP nano-column to an emitter nanospray needle for real time ionization and peptide fragmentation in an Orbitrap QExactive mass spectrometer (Thermo Fisher). During the chromatography run, we examined an enhanced FT-resolution spectrum (resolution=70,000) followed by the HCD MS/MS spectra from the 20 most intense parent ions. Dynamic exclusion was set at 40 s. For increased proteome coverage, labelled samples were also fractioned by cation exchange chromatography (Oasis HLB-MCX columns); fractions were desalted and analysed using the same system and conditions described before.

#### Protein identification and quantification

 All spectra were analysed with Proteome Discoverer (version 1.4.0.29, Thermo Fisher Scientific) using SEQUEST-HT (Thermo Fisher Scientific). The Uniprot database, containing all mouse sequences (March 03, 2013), was searched with the following parameters: trypsin digestion with 2 maximum missed cleavage sites; precursor and fragment mass tolerances of 2 Da and 0.03 Da, respectively; methionine oxidation as a dynamic modification; and carbamidomethyl cysteine and N-terminal and Lys iTRAQ modifications as fixed modifications. Peptides were identified using the probability ratio method^73^, and false discovery rate (FDR) was calculated using inverted databases and the refined method ^70^ with an additional filtering for a precursor mass tolerance of 15 ppm^74^. Identified peptides had a FDR equal to or lower than 1% FDR.

Proteins were quantified from reporter ion intensities and quantitative data analysed with QuiXoT, based on the WSPP statistical model ^69^,^70^. In this model, protein log2-ratios are expressed as standardized variables, i.e. in units of standard deviation according to their estimated variances (Zq values).

### Liquid chromatography/mass spectroscopy analysis of exosome glycerophospholipids

Exosome samples from WT and KO mice corresponding to 2.5 mg protein were used. Before lipid extraction, the following internal standards were added: 200 pmol each of 1,2- dipentadecanoyl-sn-glycero-3-phosphocholine, 1,2-dilauroyl-sn-glycero-3- phosphoethanolamine, 1,2-dipalmitoyl-sn-glycero-3-phosphoinositol, 1,2-dimiristoyl-sn-glycero-3-phosphoserine, 1,2-dimiristoyl-sn-glycero-3-phosphate, and 1-heptadecanoyl-2- arachidonoyl-3-phosphoglycerol, according to the method of Bligh and Dyer^75^. After evaporation of organic solvent under vacuum, the lipids were redissolved in 50 μl solvent mixture (75% A, 25% B) and 40 µL was injected into an Agilent 1260 Infinity high-performance liquid chromatograph equipped with an Agilent G1311C quaternary pump and an Agilent G1329B Autosampler. The column was a FORTIS HILIC (150 × 3 mm, 3 µm particle size) (Fortis Technologies) protected with a Supelguard LC-Si (20 mm × 2.1mm) guard cartridge (Sigma-Aldrich). Glycerophospholipids and sphingomyelin were separated according to the procedure described by Axelsen et al.^76^, with minor modifications. Briefly, the mobile phase consisted of a gradient of solvent A (hexanes:isopropanol, 30:40, by volume) and solvent B (hexanes:isopropanol:20 mM AcONH_4_ in water, 30:40:7 by volume). The gradient started at 75% A from 0 to 5 min, then decreased from 75% A to 40% A at 15min, from 40% A to 5% A at 20 min, holding at 5% until 40 min, and increasing to 75% at 41min. The column was then re-equilibrated by holding at 75% A for an additional 14 min before the next sample injection. The flow rate through the column was fixed at 0.4 ml/min, and this flow entered into the electrospray interface of an API2000 triple quadrupole mass spectrometer (Applied Biosystems). The source parameters were set as follows: ion spray voltage, −4500 V; curtain gas, 20 psi; nebulizer gas, 35 psi; desolvation gas, 65 psi; temperature, 400°C. Phospholipid species were analysed in Q1 with negative ionization. Compound parameters were fixed for all analysed species: declustering potential, −65 V; entrance potential, −10 V; focusing potential, −300 V; and collision cell exit potential, −10 V. Phospholipid species were detected as [M-H]^-^ ions except for phosphatidylcholine and sphingomyelin species, which were detected as [M+OAc^-^]^-^ adducts. For quantification, chromatographic peaks of each species were integrated and compared with the peak area of the internal standard corresponding to each phospholipid class. Since bis(monoacylglycerol)phosphate and sphingomyelin internal standards were unavailable at the time of analysis, these species were quantified using 1- heptadecanoyl-2-arachidonoyl-3-phosphoglycerol and 1,2-dipentadecanoyl-sn-glycero-3- phosphocholine standards, respectively.

### Subcellular fractionation

Intracellular compartments were separated by sucrose density gradient centrifugation. Cells were washed three times in 10 mM Tris-HCl pH 7.5, 0.5 mM MgCl_2_ and then scraped and resuspended in 50 mM Tris-HCl pH 7.5, 0.25M sucrose. Cells were homogenized with 30 strokes using a Dounce homogenizer. The homonogenates were then centrifuged at 3000g for 15 min at 4°C. The supernatant was placed over a continuous sucrose gradient (2M at the bottom-0.25M at the top) and samples were separated at 100,000g for 18 hours in a SW40 rotor (Beckman) at 4°C. After centrifugation, 1 ml fractions were collected from the tube, yielding a total of 12 fractions. Cold acetone (1 ml) was added to each tube and the homogeneous mixture was allowed to precipitate at −20°C for 2 h. The samples were centrifuged at 16,000g at 4°C and the protein pellets were allowed to dry for 2 h. The protein precipitates were then analysed by SDS-PAGE for proteins, including the ER marker calreticulin and the MVB marker CD63.

### Preparation of detergent-resistant membrane fractions

DRMs from exosomes derived from Cav1WT and Cav1KO MEFs were purified as described in Navarro-Lerida et al. *JCS* (2002) with the Triton X-100 concentration reduced from 1% to 0.5%.

### Transwell migration assay

The transwell migration assay was performed with Boyden chambers containing 8 µm pore-size polycarbonate filter (Corning costar). Serum-free medium (300μl) with or without exosomes was placed in the wells of a 24-well plate. The transwell inserts were placed in the wells and 5×10^4^ MDA-MB468 cells in 200 μl serum-free medium were added to the upper chamber. After incubation for 24 h, cells on the lower membrane surface were fixed and stained with crystal violet and counted under a microscope.

### Animal model

All animal procedures conformed to EU Directive 86/609/EEC and Recommendation 2007/526/EC regarding the protection of animals used for experimental and other scientific purposes, enforced in Spanish law under Real Decreto 1201/2005. Cav1WT and Cav1-/- exosomes were labelled with PKH67 and intravenously injected every day for one week into the tail vein of TnC^-/-^ mice^77^ bred into the FVB background. After 4 days, lungs and livers were collected and embedded sections were analysed by immunohistochemistry with anti-TnC antibody (Sigma).

### Immunofluorescence microscopy

Cells grown on glass coverslips were washed twice with PBS and fixed for 15 min at room temperature in 4% paraformaldehyde in PBS. Fixed cells were washed extensively with PBS and permeabilized for 5 min in PBS containing 0.1% Triton X100 and 2% BSA to reduce non-specific binding. Cells were incubate for 1 h at room temperature in PBS containing 0.2% BSA with primary antibodies (typically diluted 1:200). Subsequent washes and incubations with Alexa fluor-647 or 546 phalloidin or fluorescent secondary antibodies were carried out in PBS for 1 h at room temperature. Coverslips were mounted in Permafluor aqueous mounting medium and examined with a Leica SPE, SP5 or SP8 confocal microscope.

### Filipin staining and quantification

Cells were grown on glass coverslips. After fixing for 10 min at 37°C with 4% paraformaldehyde in phosphate-buffered saline (PBS), cells were permeabilized with 0.2% BSA and 0.1% saponin in PBS (blocking solution) for 30 minutes. Samples were then incubated with primary antibody (LBPA) and fluorescently labeled-conjugated secondary antibodies for 1h each, interspersed with three washes in blocking solution. After an additional wash in PBS, cells were incubated for 1h with filipin (50µg/ml in PBS). After three washes in PBS, slides were mounted in Fluoromount. Fiji software was used to quantify filipin fluorescence intensity (http://fiji.sc/). Cells were segmented based on LBPA staining. Filipin staining intensity in LBPA-positive vesicles was measured after intensity thresholding.

### Quantitative real-time PCR

Frozen tissues or cell lines were analysed for specific gene expression using SybrGreen PCR Reagents (Applied Biosystems) and specific primers: mouse Tsg101 sense 5’- TCTAACCGTCCGTCAAACTGT-3’, antisense 5’-TTGTACCAGTGAGGTTCACCA-3’; mouse nSMase 2 sense 5’-ACACGACCCCCTTTCCTAATA-3’, antisense 5’- GGCGCTTCTCATAGGTGGTG-3’; mouse nSMase 1 sense 5’- TGGGACATCCCCTACCTGAG-3’, antisense 5’-TAGGTGAGCGATAGCCTTTGC-3’. Total RNA was extracted from tissues or cells using the RNeasy kit (Qiagen) and reverse-transcribed. Quantitative real-time PCR was performed with a 7500Fast Real Time PCR System. Relative expression was normalized to GADPH and HPRT.

### Immunohistochemical staining

Harvested tissues were fixed in 4% paraformaldehyde, processed through graded sucrose, embedded in OCT medium (Tissue-Tek) and stored at −80°C. Frozen sections (20 µm) were dried for 10 minutes at room temperature and blocked for 1h in PBS containing 4% chicken serum and 0.3% TritonX-100. Sections were stained with anti-TnC antibody diluted 1:200 in blocking solution overnight at 4°C. Samples were washed extensively in PBS, 0.15% TritonX- 100 for at least for 3 h. Samples were then incubated overnight with anti-rabbit Alexa fluor- 647 secondary antibody (1:200), Alexa-fluor 546-phalloidin (1:500) and Hoescht 33342 in blocking solution. After extensive washing, sections were mounted in Permafluor mounting medium. Confocal Z-stack images were captured and TnC deposits were quantified.

### Extracellular TnC and FN fiber quantification

The procedure for image processing and quantification of “fibreness” was developed as a macro in Fiji (ImageJ 1.50e x64). Fibreness measures the amount of fibre-like structures in an image, providing a readout that takes into account both the density of fibres and their length, independently of orientation. Noise reduction first stabilizes the image background. The structural information from the eigen values of the Hessian matrix is then used to apply a Frangi vesselness filter^74^ to enhance very thin, almost unidimensional tubular structures (filter scale: s=0.179 µm, close to pixel size). The output is a fibre-enhanced image where each pixel contains a fibreness score. M0 readout was computed as the mean fibreness score in the whole image. The processing pipeline and extracted M0 measures are illustrated in Supplementary Figure 3b.

### Statistical analysis

Error bars depict SEM. Statistical significance was determined with GraphPad Prism by unpaired Student’s *t*-test or Mann-Whitney test, as indicated; * p<0.05, ** p<0.01, *** p<0.001.

## ACKNOWLEGMENTS

We thank the CNIC Microscopy and Histology Units for technical assistance. We thank Daniel Jiménez-Carretero of the CNIC Cellomics Unit for developing the software for TnC fibre quantification. We also thank Dr. Kirchner for providing plasmids encoding ubiquitinated Cav1 mutants, Fátima Sánchez-Cabo of the CNIC Bioinformatics Unit for help with the statistical analysis of exosome populations, Dr. Germán Andrés for help with electron microscopy, and Dr. Fidel Lolo for PTRF-reconstituted PTRFKO cells. This study was supported by grants to M.A.d.P. from the Ministerio de Ciencia, Innovación y Universidades (MICINN: SAF2011- 25047, CSD2009-0016, SAF2014-51876-R, SAF2017-83130-, the Fundació La Marató de TV3 (674/C/2013), the Worldwide Cancer Research Foundation (AICR 15-0404) and Fondo Europeo de Desarrollo Regional (FEDER) *“Una manera de hacer Europa”*. M.A.d.P´s group received funding from the European Union Horizon 2020 research and innovation programme under Marie Sklodowska-Curie grant agreement n° 641639. J. B. was supported by MICINN grants SAF2013-48201-R and SAF2016-80883-R, and G. O. was supported by INSERM, the University of Strasbourg, the Ligue contre le Cancer, and the Institut National du Cancer (INCa). LA-A was supported by a MICINN predoctoral fellowship associated with the Severo Ochoa Excellence program (FPI). IN-L was supported by a postdoctoral fellowship from the Asociación Española Contra el Cáncer (AECC). Miguel Sánchez Àlvarez (CNIC) provided help with the manuscript preparation. Simon Bartlett (CNIC) provided English editing. The CNIC is supported by the Ministerio de Ciencia, Innovación y Universidades and the Pro CNIC Foundation, and is a Severo Ochoa Center of Excellence (SEV-2015-0505)

## Abbreviations

Extracellular matrix: (ECM)
Caveolin-1: (Cav1)
Tenascin-C: (TnC)
Cancer- Associated-Fibroblast: (CAF)
Mouse embryonic fibroblast: (MEF)
Multivesicular body: (MVB)
Lyso-bis-phosphatidic acid: (LBPA)
Intraluminal vesicle: (ILV)
Neutral sphingomyelinase-2: (nSMase-2)
Endoplasmic reticulum: (ER)
Knock out: (KO)
Knock down: (KD)
Epithelial- mesenchymal transition: (EMT)
Polymerase I and transcript release factor: (PTRF)

